# Ibotenic acid biosynthesis in the fly agaric is initiated by glutamate hydroxylation

**DOI:** 10.1101/846857

**Authors:** Sebastian Obermaier, Michael Müller

## Abstract

The fly agaric, *Amanita muscaria*, is widely known for its content of the psychoactive metabolites ibotenic acid and muscimol. 50 years ago, their biosynthesis was hypothesized to start with 3-hydroxyglutamate. Here, we build on this hypothesis by the identification and recombinant production of a glutamate hydroxylase from *A. muscaria*. The corresponding gene is surrounded by six other genes, which we link to ibotenic acid production using recent genetic data. Our data provide new insights into a decades-old question concerning a centuries-old drug.

## Main

*Amanita muscaria*, the fly agaric, is perhaps the most prominent of all mushrooms, known for its extravagant appearance with a red cap covered by white specks, and its infamous toxicity. The primary bioactive compounds of *A. muscaria* are ibotenic acid and its decarboxylation product muscimol (1–3). Although actually less toxic than generally assumed, the psychoactive effects of ibotenic acid and muscimol derive from their structural similarity to the neurotransmitters glutamate and GABA, respectively (4, 5) (Figure 1A).

**Figure 1.**
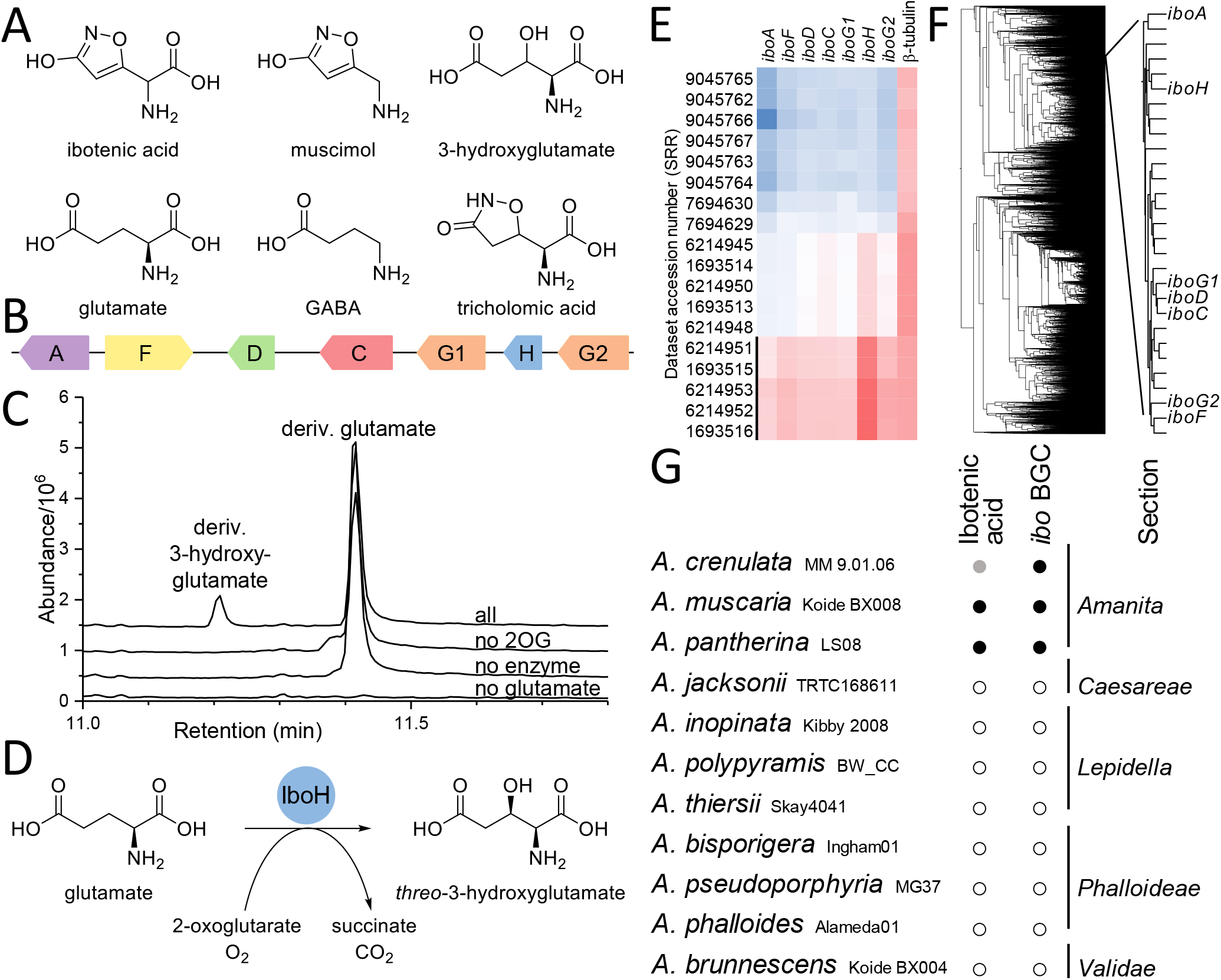
The ibo biosynthetic gene cluster. (A) Structures of A. muscaria metabolites and analogs. (B) Schematic of the ibo BGC. (C) GC-MS total ion chromatograms of IboH assays. 3-Hydroxyglutamate formation was dependent on 2-oxoglutarate (2OG), enzyme, and glutamate. (D) Stereoselective glutamate hydroxylation by IboH. (E) Normalized expression of ibo genes across RNA-seq datasets from NCBI SRA: low expression (blue) to high expression (red). The black line marks cocultivation experiments of A. muscaria with Populus tremula x tremuloides. The β-tubulin gene is included for comparison. (F) Unbiased coexpression clustering of 11915 expressed A. muscaria genes. The ibo genes cluster closely together, indicating coregulation. (G) Presence of the ibo genes and (putative) ibotenic acid production of Amanita species with sequenced genomes/transcriptomes. The three strains containing ibo genes belong to Amanita section Amanita.

Shortly after the structure of ibotenic acid was solved in 1964, hypotheses about its origin in fungi were made (1). Eugster and coworkers hypothesized that ibotenic acid is derived from 3-hydroxyglutamate (6), which would necessitate an enzyme that catalyzes the hydroxylation of glutamate. However, the biosynthesis of ibotenic acid and muscimol has remained obscure.

To identify the biosynthetic genes, we assumed that the formation of ibotenic acid is initiated with the hydroxylation of either glutamine or glutamate. Hitherto, no enzyme that catalyzes this reaction on free substrate has been experimentally verified (7). However, 3-hydroxyglutamine occurs as a component of the antifungal pneumocandin B_0_ from *Glarea lozoyensis*. Its biosynthetic gene cluster (BGC) includes a putative dioxygenase, GloE, which has been proposed as a candidate enzyme for the hydroxylation of glutamine (8). Therefore, we used its protein sequence to screen the *A. muscaria* genome (9). Indeed, a homologous protein, IboH (KIL56739), is encoded in a genetic region that resembles a BGC. This putative *ibo* BGC (Figure 1B) comprises a cytochrome P450 monooxygenase (IboC KIL56737), a flavin-dependent monooxygenase (IboF KIL56733), an adenylating enzyme (IboA KIL56732), two similar pyridoxal phosphate-dependent enzymes (IboG1 KIL56738 and IboG2 KIL56740), and a decarboxylase (IboD KIL56734). The genes include all functionalities theoretically needed for the biosynthesis of ibotenic acid (see below).

Consequently, the assignment of the BGC was verified experimentally. The *iboH* gene was expressed in *Escherichia coli* with an N-terminal GST tag (MN520442). As IboH is predicted to be an Fe(II)/2-oxoglutarate-dependent dioxygenase, the purified protein was incubated aerobically with Fe^2+^, 2-oxoglutarate, ascorbic acid, and the putative substrate glutamine or glutamate. The reaction mixtures were derivatized with ethyl chloroformate/ethanol and analyzed by GC-MS. While the glutamine assay was negative, glutamate was transformed to a product that was detected as a new peak in the GC-MS chromatogram (Figure 1C, 11.2 min). Control experiments lacking either enzyme, glutamate, or 2-oxoglutarate did not yield the product. As the yield with the purified enzyme was too low for isolation, a whole-cell approach was used: incubation of live *E. coli GST-iboH* cells with L-glutamate gave sufficient amounts of the product, which was extracted from the cell supernatant by cation exchange chromatography as the hydrochloride. NMR analysis showed the presence of *threo*-3-hydroxyglutamate^1^, confirming the role of IboH to be that of an L-glutamate 3-(*R*)-hydroxylase (Figure 1D).

To determine the biological relevance in *A. muscaria*, mushroom samples (collected near Feldberg, Black Forest, Germany) were analyzed by GC-MS. Ibotenic acid was detected along with low levels of 3-hydroxyglutamate. This hints towards IboH being active in its native organism, coinciding with the production of ibotenic acid. Furthermore, public RNA-seq data (10) revealed that the seven genes in the *ibo* BGC are highly expressed when *A. muscaria* was artificially grown in symbiosis with *Populus*, which is close to its natural lifestyle (Figure 1E). To investigate whether the genes are functionally linked, coexpression network analysis was conducted. The data showed that the genes have a very similar expression pattern, which indicates coregulation and hence a common metabolic function (11) (Figure 1F).

To further probe the link of the identified BGC to the production of ibotenic acid, other *Amanita* species were searched for homologous genes. *Amanita pantherina* is a verified producer of ibotenic acid (12), and reports suggest that ingestion of *Amanita crenulata* can cause symptoms that resemble those of ibotenic acid and muscimol (13). As transcriptome data were available for these two species (14, 15), the RNA-seq reads were mapped to the *ibo* BGC. Indeed, the data demonstrate that both species actively express close homologs of each of the *ibo* genes (> 80% nucleotide identity). Additionally, we analyzed genomes of eight *Amanita* species from four taxonomic sections, which do not produce ibotenic acid. None of these contain the *ibo* BGC, substantiating the correlation between ibotenic acid and the *ibo* genes (Figure 1G). Apparently, the presence of the *ibo* BGC is confined to *Amanita* section *Amanita*, which is in accordance with previous studies on the taxonomic distribution of ibotenic acid production (16, 17).

To deduce the biosynthetic functions of the *ibo* proteins, their sequences were checked against known enzymes. From this, we propose the functions illustrated in Figure 2 and described in the following. Similar proteins are listed in square brackets along with sequence identity values for comparison.

**Figure 2.**
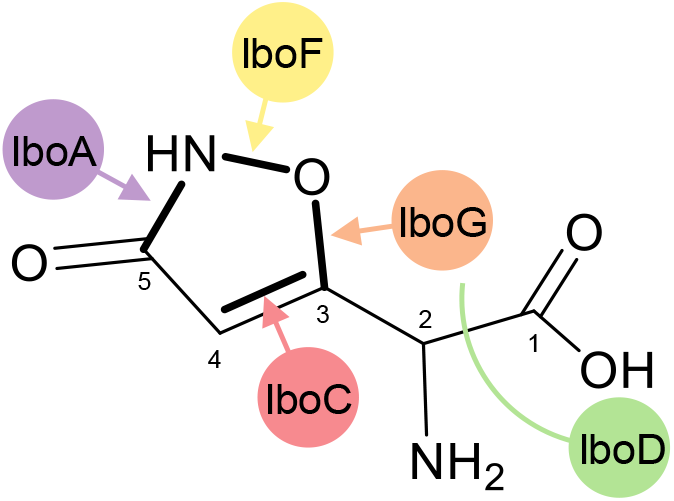
Structure of ibotenic acid and the proposed roles of the identified enzymes in its biosynthesis. Starting from 3-hydroxyglutamate, newly formed bonds are highlighted in bold.

The first committed step is glutamate hydroxylation by IboH, and the last step is likely decarboxylation (18) of ibotenic acid to muscimol by IboD [tryptophan decarboxylase (19) P0DPA6, 32%]. The order of the intermediate reactions is somewhat ambiguous. IboA [adenylate-forming reductase (20) F8P9P5, 21%] likely activates the terminal carboxylic acid function to introduce an amide bond. The flavin monooxygenase IboF [B8NM63, B8NM73, 21–24%, (21)] likely introduces the N–O bond. There are two options for this step. Either IboF directly hydroxylates the amide nitrogen formed by IboA, or it hydroxylates an external N-containing compound, whose N–O bond is subsequently introduced [cf. cycloserine (22) and trichostatin (23) biosynthesis]. The paralogous pyridoxal-dependent enzymes IboG1 and IboG2 [I1RZK8, 38–39%, (24)] are likely involved in substitution of the OH group at position 3 by the O–N moiety. While the order of these reactions cannot be inferred with certainty, the first cyclic intermediate is probably tricholomic acid (25) (a metabolite found in *Tricholoma muscarium*). Desaturation of tricholomic to ibotenic acid is likely catalyzed by the cytochrome P450 IboC [A1CFL5, A1CFL6, A0A286LF02, 27–30% (19, 26)].

Taken together, the findings described above indicate that the *ibo* genes are responsible for ibotenic acid production in at least three *Amanita* species. The identified BGC contains the glutamate hydroxylase IboH, whose activity was demonstrated in a heterologous system. This discovery revives the long-dormant research question on toxification of the fly agaric.

## Methods

### GC-MS analysis

3-Hydroxyglutamate derivatization, in a 2 mL microcentrifuge tube: 100 μL sample, 50 μL ethanol, and 100 μL 8% ethyl chloroformate in dichloromethane were shaken for 15 s. 20 μL pyridine was added and the mixture was shaken with an open lid for 45 s. 250 μL ethyl acetate was added and the tube was shaken for 1 min. The upper layer was concentrated to dryness under reduced pressure, then dissolved in 40 μL ethyl acetate. Ibotenic acid derivatization was done as described (27). System: Agilent HP 6890N GC, HP 5973 mass detector, DB-5ms column (30 m, 0.25 mm, 0.25 μm), helium 1.0 mL/min, 60 °C 0–3 min, to 280 °C 3–14 min, 280 °C 14–19 min.

### IboH conversion of glutamate

Glutathione S-transferase-tagged protein (GST-IboH) was purified from *E. coli* BL21-Gold (DE3) pGEX-6P-1-*iboH* cells using glutathione affinity chromatography. Assays in PBS pH 7.2, 2.4 mM ascorbic acid, 0.18 mM FeSO_4_, 8 mM sodium 2-oxoglutarate, 3.2 mM sodium glutamate, 2.3 mg/mL enzyme. For whole-cell conversion, the same strain was grown at 23 °C in LB medium with 0.29 mM ampicillin, 0.72 mM FeSO_4_, 5.3 mM sodium glutamate; induction at OD_600_ 1.0 with 0.4 mM IPTG. After 24 h, cells were transferred to minimal medium with ampicillin, IPTG, 53 mM sodium glutamate, 2% glucose; incubation for 20 h at 23 °C. Cell supernatant (75 mL) was adjusted to pH 2 with HCl, applied to a Dowex 50WX8 column (H^+^ form, 200–400 mesh, 10 × 153 mm), washed with water, and eluted with 0.07 mM HCl. Retention volume was 9.1 column volumes [cf. published values (28, 29)]. The product* was precipitated from methanol/acetonitrile. ^1^H NMR (400 MHz, D_2_O, internal acetonitrile δ_CH3_ 2.06 ppm), δ = 2.72 (dd, *J* = 16.4, 8.8 Hz, 1H), 2.87 (dd, *J* = 16.4, 4.3 Hz, 1H), 4.09 (d, *J* = 4.8 Hz, 1H), 4.62 (ddd, *J* = 8.8, 4.8, 4.3 Hz, 1H). GC-MS 11.2 min, *m*/*z* (%) = 273 (1, [M–H_2_O]^+^), 228 (11), 200 (55), 182 (17), 172 (59), 156 (33), 155 (41), 154 (48), 144 (21), 128 (100), 127 (59), 126 (79), 99 (58).

### RNA-seq analysis

RNA-seq data analysis with the Galaxy platform. Short read mapping with HISAT 2.1.0. *A. muscaria*: default settings; *A. crenulata* (SRR485870) and *A. pantherina* (SRR6823443–SRR6823454): coefficients for the alignment score function [f(x) = B + A · x], B = 0, A = −0.9; maximum intron length = 1000.

### Coexpression analysis

Calculation of counts with StringTie 1.3.4d, using NCBI-SRA transcriptome datasets given in Figure 1E. Normalization with limma-voom 3.38.3 (filter lowly expressed genes 4 CPM/4 samples). In R/RStudio: construction of the coexpression network with WGCNA (30). Coexpression analysis was independent from gene location in the genome. Dendrogram in Figure 1F with FigTree 1.4.3. For expression matrix and R script, see supporting information.

## Supporting information

Amanita muscaria expression count matrix

R script for coexpression network analysis

## Footnotes

1 *threo*-3-Hydroxyglutamate occurred along with various amounts of a byproduct (2–9-fold excess), which coeluted on ion-exchange chromatography. 2D NMR indicated the byproduct has the same carbon chain as 3-hydroxyglutamate, and it probably originates from chemical modification by the *E. coli* host, e.g. formation of a dimeric ester. ^1^H NMR (400 MHz, D_2_O), δ = 2.85 (dd, *J* = 17.9, 4.2 Hz, 1H), 2.93 (dd, *J* = 17.9, 8.9 Hz, 1H), 4.31 (ddd, *J* = 8.9, 4.2, 3.5 Hz, 1H), 4.79 (d, *J* = 3.5 Hz, 1H). GC-MS after derivatization showed only the major peak at 11.2 min (Figure 1C).

